# Effects of Face Repetition on Ventral Visual Stream Connectivity using Dynamic Causal Modelling of fMRI data

**DOI:** 10.1101/2022.07.05.498907

**Authors:** Sung-Mu Lee, Roni Tibon, Peter Zeidman, Pranay S. Yadav, Richard Henson

## Abstract

Stimulus repetition normally causes reduced neural activity in brain regions that process that stimulus. Some theories claim that this “repetition suppression” reflects local mechanisms such as neuronal fatigue or sharpening within a region, whereas other theories claim that it results from changed connectivity between regions, following changes in synchrony or top-down predictions. In this study, we applied dynamic causal modelling (DCM) on a public fMRI dataset involving repeated presentations of faces and scrambled faces to test whether repetition affected local (self-connections) and/or between-region connectivity in left and right early visual cortex (EVC), occipital face area (OFA) and fusiform face area (FFA). Face “perception” (faces versus scrambled faces) modulated nearly all connections, within and between regions, including direct connections from EVC to FFA, supporting a non-hierarchical view of face processing. Face “recognition” (familiar versus unfamiliar faces) modulated connections between EVC and OFA/FFA, particularly in the left hemisphere. Most importantly, immediate and delayed repetition of stimuli were also best captured by modulations of connections between EVC and OFA/FFA, but not self-connections of OFA/FFA, consistent with synchronization or predictive coding theories.

## 1 Introduction

Repetition suppression (RS) refers to decreased neural responses produced by repeated exposures to stimuli. RS is observed in human studies using functional magnetic resonance imaging (fMRI; for review, see Grill-Spector et al., 2006) and has been associated with the behavioural phenomenon of “priming”, i.e., faster and/or more accurate responses to repeated stimuli. RS has also been used as a tool to infer functional characteristics of neural populations, particularly in sensory regions (also called “fMRI adaptation”, Grill-Spector & Malach, 2001; Larsson et al., 2016). For example, two regions in the ventral visual stream - the bilateral fusiform face area (FFA) and the occipital face area (OFA) - consistently show RS to repeated presentations of the same faces (Henson, 2016).

### 1.1 Neural theories of Repetition Suppression

As described below, four main theories have been developed to account for RS: Fatigue, Sharpening, Synchronization and Predictive coding (Grill-Spector et al., 2006; Wiggs & Martin, 1998; Gotts et al., 2012; Henson, 2016). Although these theories have focused on different features of repetition effects, the aim of the present study was to test their predictions in terms of effective connectivity between face-responsive regions. To this end, we applied dynamic causal modeling (DCM, Friston et al., 2003) on a publically available fMRI dataset that includes initial and repeated presentations of familiar, unfamiliar and scrambled faces (Wakeman & Henson, 2015). DCM is a Bayesian framework for comparing models with specified connectivity within a network of regions of interest (ROIs). It incorporates a generative model of fMRI data, in which connections are represented by three ROI-by-ROI matrices of parameters: the A matrix represents the fixed (or endogenous) directional connections from one ROI to another; one or more B matrices represent the modulation of the corresponding endogenous connections due to one or more experimental manipulations (e.g., each type of stimulus); and the C matrix is the direct (or exogenous) input to one or more ROIs for one or more experimental manipulations. The dynamics (fMRI timeseries) of the ROIs are then modelled using 1) a parameterized differential equation that expresses the rate of change of neural activity in each ROI as a function of the level of activity in every connected ROI, triggered by the timing of each exogenous input; and 2) a haemodynamic model that transforms the predicted neural activity into the fMRI BOLD signal, with a small number of haemodynamic parameters than can vary across ROIs. When applying DCM to RS paradigms, the C matrix can code all stimuli, whereas the B matrix can code for the difference between initial and repeated presentations of stimuli (for further details about DCM, see Methods).

Of the four theories of RS mentioned above, the first two emphasize changes within an ROI, which is captured in DCM by the self-connections (diagonal terms of A and B matrices). These are constrained to be negative to ensure the network dynamics are stable (i.e., activity eventually returns to zero at some time after an exogenous input). According to the Fatigue theory (Li et al., 1993; McMahon & Olson, 2007), it is the neurons that are most selective for a stimulus (and therefore show the greatest firing to that stimulus) that show greatest reduction in firing when that stimulus is repeated. This pattern has been observed in early visual cortex (Avidan et al., 2002), and may underlie some forms of perceptual adaptation (e.g., Grill-Spector et al., 1999). The mechanism underlying the fatigue could be firing-rate adaptation or synaptic depression (Grill-Spector et al., 2006). If these mechanisms operate primarily within an ROI (see Discussion), then in the DCM framework, they would be captured by changes in the self-connections.

The Sharpening theory (Wiggs & Martin, 1998) also offers a “within-region” perspective, but makes the opposite neural predictions to fatigue theory: i.e., it is the less selective neurons whose firing is reduced (as opposed to the most selective), resulting in a sparser distribution of firing. The mechanisms of sharpening might include strengthening of inhibitory, lateral connections between neurons. Consistent with the sharpening model, Jiang et al. (2006, 2007) found that perceptual training sharpens the tuning of neurons. At a population-level measured by fMRI (i.e., when averaging over many neurons within an ROI), because there tend to be more non-selective than selective neurons, the mean firing rate decreases (causing RS). This would again be apparent in DCM by changes in the self-connections of the B matrix, e.g., increased self-inhibition, indistinguishable from Fatigue theory.

The remaining two theories assume that RS also arises from connections between regions. The Synchronization theory of Gotts et al. (2012) proposes that repetition leads to increased synchronization of neural activity across regions, such that greater communication can be achieved despite lower mean firing rates. The reduced firing rates cause RS, while the increased synchrony causes more efficient neural processing and hence behavioural effects like priming (Ghuman et al., 2008). This increased neural synchrony is likely to be associated with stronger effective connectivity between regions, corresponding to off-diagonal elements in DCM’s B matrix.

Finally, the Predictive Coding theory (Friston, 2005; Henson, 2016) suggests that RS is associated with changes in effective connectivity between regions, as well as within a region. This theory proposes that neurons at one level of a hierarchy receive predictions from higher levels, and feed forward the difference (i.e., prediction error) between these predictions and the input from levels below. When a stimulus has been processed before, the predictions are improved, and therefore the prediction error is reduced. A single ROI (as resolved by standard fMRI) is assumed to contain neurons receiving predictions from the level above, neurons receiving prediction errors from the level below, and neurons sending the resulting prediction error to the level above (even if these neurons are in different layers of cortex; Friston, 2005). However, assuming that the fMRI signal is dominated by the feedforward neurons whose firing codes prediction error (Egner et al., 2010), the mean fMRI response will be reduced by repetition. Though the mapping from neural interactions to fMRI effective connectivity is not simple (see Discussion), the improved predictions might be expected to affect backward connections in DCM (e.g., from FFA to OFA), while the reduced prediction errors fed forward might be expected to affect forward connections (e.g., from OFA to FFA).

A previous study of Ewbank et al. (2013) used DCM to investigate effects of face repetition (within and across changes in the size of images), and found evidence that repetition modulated connections from right OFA to right FFA, supporting the synchronization/predictive coding account. However, the RS data in that study came from a blocked fMRI design, comparing blocks in which the same face was shown multiple times against blocks in which a new face was shown in each trial. This blocking means that participants can expect whether or not the next stimulus is a repeat, which is also known to reduce the fMRI response (Summerfield et al., 2008; also called “expectation suppression”, Grotheer & Kovács, 2015). In the present data, initial and repeated trials were pseudo-randomly intermixed, dramatically reducing the ability to accurately expect the next stimulus type. Furthermore, the repetition in the Ewbank et al.’s DCM studies was always immediate (i.e., no intervening stimulus), yet the lag between initial and repeated presentations may affect the mechanisms of RS (e.g., immediate repetition may engage fatigue to a greater extent than longer-lag repetition; see Epstein et al., 2008; Henson, 2016). In the present data, there were also delayed repetitions (with several intervening faces), which allowed testing of whether the effects of repetition on connectivity differ by repetition lag. Finally, Ewbank et al. (2013) only included 2 ROIs in their DCM: the OFA and FFA in the right hemisphere. While this allowed testing of whether repetition affects forward, backward and/or self-connections, it did not allow for the possibility that repetition already affects the input to these regions, e.g., in the forward connectivity from earlier regions in a processing stream, such as early visual cortex (EVC), which limited the ability to test some specific theories of face processing (see below). Moreover, because the DCM only included ROIs in the right hemisphere, it did not allow testing of whether the same repetition effects occur in the left hemisphere. Notably, though face-related OFA and FFA activations in fMRI are often stronger/more selective in the right hemisphere, paralleling suggestions from brain lesions that the right hemisphere is specialized for face processing (Ishai et al., 2005; Rossion, 2018), similar face-related activations are found in the left hemisphere. We addressed these issues by comparing a “2-ROI” network in each hemisphere separately (as in Ewbank et al., 2013), with a “3-ROI” network that also included EVC in each hemisphere, and a “6-ROI” network that included bilateral EVC, OFA and FFA ROIs.

### 1.2 Network theories of Face Processing

Finally, though we have focused on repetition effects, our DCM models also allowed testing of other hypotheses associated with face processing. Firstly, we have assumed above (e.g., in the discussion of forward and backward connections according to the Predictive Coding theory) that FFA sits “higher” than the OFA in a face processing hierarchy, as common in theories of face processing (Haxby et al., 2000; Fairhall & Ishai, 2007). However, Rossion and colleagues have suggested that information may flow from EVC to FFA first, and then back from FFA to OFA. This is based on neuroimaging findings from patients with OFA lesions, who still show face-related activation in FFA in the same hemisphere (Rossion et al., 2003; Gentile et al., 2017; Steeves et al., 2009). This suggests a direct connection from EVC to FFA that does not go via OFA (or else input from the contralateral FFA), meaning there is not a strict, sequential hierarchy from EVC to OFA to FFA.

The first DCM study of face processing (Fairhall & Ishai, 2007) found that face perception modulated connections from OFA to FFA, favouring the conventional feedforward, hierarchical view. A more recent meta-analysis of four DCM fMRI experiments (Kessler et al., 2021) replicated the increased “forward” connectivity from OFA to FFA for faces, but also found more negative “backward” connectivity from FFA to OFA. However, neither of these studies allowed for input from EVC to both ROIs, which would allow, for example, face-related input to FFA that bypasses OFA, as suggested by the patient fMRI data of Rossion and colleagues. Nor did either study allow modulations of self-connections by faces, nor modulation of connections between the two hemispheres. A study by Frässle et al. (2016) tested DCMs that connected OFA and FFA in both hemispheres, as well as input from left and right EVC. Their findings highlighted the role of interhemispheric integration between bilateral OFA in face perception, in addition to feedforward modulations from EVC to OFA and then FFA. However, they did not allow the direct connections between EVC and FFA suggested by Rossion and colleagues (nor allow modulation of self-connections). We therefore revisited this question in the present dataset, operationalizing face “perception” by contrasting faces with scrambled faces, and adding another “B” matrix to capture modulation of connections by face perception. By allowing the connections from EVC to be modulated by both faces and by repetition, we could test alternative hypotheses that face-related activation and RS respectively already arrive in the input to OFA and/or FFA, through altered synaptic weights from earlier visual regions.

Finally, there is also debate around the role of FFA and OFA in face recognition, as operationalized in the present dataset by contrasting familiar faces (known to participants) with unfamiliar faces (see also Henson et al., 2003). While impairments of face recognition/identification, despite intact face perception, are more often associated with lesions to more anterior temporal lobe (ATL) regions (Damasio et al., 1990; Gainotti & Marra, 2011), neuroimaging studies sometimes show additional activation of FFA for familiar than unfamiliar faces (Henson et al., 2003). While this familiarity-related activation could reflect feedback from ATL, we did not include an ATL ROI in the present DCMs because no such region showed any differential activity in the whole-brain contrasts, possibly because of susceptibility-related fMRI signal dropout in this dataset. Nonetheless, we could at least test whether familiarity effects reflected local effects (self-connections), connectivity from OFA to FFA, or between left and right hemispheres for example.

### 1.3 Methodological Advancements

The previous meta-analysis by Kessler et al. (2021) made inferences about individual parameters (connections) in their DCM models of a face network. However, this parameter-level inference ignores potential covariances between the posterior estimates of those parameters, which limits their reproducibility and interpretability (Rowe et al., 2010). Here, we focused on model-level inference, which accommodates covariances between all parameters. To address our hypotheses, we performed binary comparisons of two families of models that differed in a certain type of connection, such as models with versus without modulation of self-connections by repetition, for example, or models with versus without modulation of forward connections from ECV to FFA by face perception. More specifically, we compared families in terms of the free energy approximation to their model evidences, converted to a posterior probability of one family being more likely than the other (where a probability of 95% was taken to be sufficient evidence to favour one family).

Furthermore, we employed recent developments in group DCM modelling, using a Parametric Empirical Bayes (PEB) approach (Friston et al., 2015). By creating a hierarchical model, empirical priors at the group level shrink the parameter estimates for individual participants toward those associated with the global maximum of the model evidence. This finesses problems due to local minima inherent in the inversion of nonlinear and ill-posed DCM models, thereby providing more robust and efficient estimates.

To summarise, the main purpose of this study was to examine critical hypotheses in terms of connectivity arising from the four theories of RS: fatigue, sharpening, synchronization and predictive coding. If repetition modulates within-region connectivity (self-modulations in DCM’s B matrix), this is consistent with the fatigue and sharpening models. If repetition modulates between-region connections (forward or backward modulations in DCM’s B matrix), this is consistent with synchronization and predictive coding models. Note also that these theories are not mutually exclusive - for instance, predictive coding may induce neural sharpening, causing changes both between and within regions. Furthermore, we took the opportunity to revisit questions about the functional architecture of face perception and face recognition, given that the dataset included repetition of unfamiliar faces, famous faces and scrambled faces.

## 2 Materials and Methods

### 2.1 Dataset

The multi-modal (MRI, EEG, MEG) human neuroimaging dataset is available on OpenfMRI (https://www.openfmri.org/dataset/ds000117/; Wakeman & Henson, 2015). It consists of 19 participants with an age range of 23–37 years (note that this is a superset of the participants available on OpenNeuro, https://openneuro.org/datasets/ds000117). Eighteen participants were included after removing one participant whose debriefing showed they did not recognize any famous faces in one run.

During each of the 9 runs, participants made left-right symmetry judgments to randomly presented images of 16 unique faces from famous people, 16 unique faces from nonfamous people (unfamiliar to participant), and 16 phase-scrambled versions of the faces. Half of the stimuli repeated immediately, and the other half repeated after delays of 5-15 stimuli intervals. We used debriefing data to re-define familiarity of each face for each participant (i.e., reclassifying famous faces they did not know as “unfamiliar” and reclassifying nonfamous faces they said they knew as “familiar”, even though latter was rare).

The MRI data were acquired with a 3T Siemens Tim-Trio MRI scanner (Siemens, Erlangen, Germany). The fMRI data came from a gradient echo-planar imaging (EPI) sequence, with TR of 2000 ms, TE of 30 ms and flip angle of 78°. A T1-weighted structural image of 1 × 1 × 1 mm resolution was also acquired using a MPRAGE sequence (for more details, see Wakeman & Henson, 2015).

### 2.2 fMRI analysis and ROI selection

The fMRI data were pre-processed using the SPM12 software (www.fil.ion.ucl.ac.uk/spm). The first two scans were removed from each session to allow for T1 saturation effects. The Matlab scripts used for all analyses that follow are available here: https://github.com/SMScottLee/Face_DCM_fMRI. The functional data were realigned to correct for head motion, interpolated across time to the middle slice to correct for the different slice times, and coregistered with the structural image. The structural image was segmented and normalized to a standard MNI template, and the normalization warps were then applied to the functional images, resulting in voxel sizes of 3 × 3 × 3 mm. These were finally spatially smoothed using a Gaussian filter of 8 mm FWHM for mass univariate statistics.

After preprocessing, fMRI data were analyzed in a two-stage approximation to a mixed-effects model. In the first stage, neural activity was modeled by a delta function at stimulus onset. The BOLD response was modeled by a convolution of these delta functions by a canonical haemodynamic response function. The resulting time courses were downsampled at the midpoint of each scan to form regressors in a General Linear Model (GLM). The experiment crossed two factors. The first factor was repetition (initial stimulus, immediately repeated or delayed repeated) and the second factor was the type of stimulus (familiar face, unfamiliar face or scrambled face). Each test session therefore contained 9 regressors of interest: initial familiar face, immediately repeated familiar face, delayed repeated familiar face, initial unfamiliar face, immediately repeated unfamiliar face, delayed repeated unfamiliar face, initial scrambled face, immediately repeated scrambled face, and delayed repeated scrambled face. To capture face processing effects and to guide ROI selection (see below), 4 contrasts were predefined: “face perception” was operationalised by contrasting all faces versus scrambled faces; “immediate repetition” was operationalised by contrasting all initial presentations (faces and scrambled faces) versus immediate repeats; “delayed repetition” was operationalised by contrasting all initial presentations versus delayed repeats; “face recognition” was operationalised by contrasting all familiar faces versus unfamiliar faces. In addition, 5 interactions between these contrasts were tested (e.g., whether immediate repetition effects were bigger for faces than scrambled faces).

The group-level, family-wise error-corrected (*p* < .05) results showed greater BOLD response to faces than scrambled faces (averaged across initial and repeated presentations) in bilateral occipital face area (OFA) and fusiform face area (FFA), plus a cluster in left orbitofrontal cortex.

While superior temporal sulcus (STS) is often associated with face processing (Babo-Rebelo et al., 2022; Haxby et al., 2000) and has been included in some previous DCM analyses (Fairhall & Ishai, 2007; Kessler et al., 2021), we did not find it in our group-level univariate results and so did not include it in the present analysis. However it should be kept in mind that some of the present effects in OFA and/or FFA could emerge from interactions with STS (or other regions like ATL; see Introduction), which could be investigated in future studies.

To allow for some individual variability in the location of OFA and FFA, and maximize their SNR, these ROIs were defined by the contrast of faces > scrambled, uncorrected *p* < .05, for each subject, but masked with a 10-mm sphere located at the group-result peak (right OFA [x = +39, y = -82, z = -10], left OFA [x = -36, y = -85, z = -13], right FFA [x = +42, y = -46, z = -19], left FFA [x = -39, y = -49, z = -22]) in order to constrain within anatomically-similar areas (Supplementary Figure 1). Because the contrast of all trials > baseline activated most of the occipitotemporal cortex (and the stimuli straddled both visual hemifields), the present data did not enable selection of distinct clusters for EVC. Therefore, ROIs for right and left EVC were defined by the contrast of left > right and right > left hemifield input from a previous study (Henson & Mouchlianitis, 2007), and masked with subject-specific contrasts of all trials > baseline in the present data, again to maximize SNR. The number of voxels per ROI ranged across participants from 140-178 for rEVC, 47-151 for rOFA and 5-157 for rFFA; 86-112 for lEVC, 12-143 for lOFA and 3-167 for lFFA.

The first singular vector of the fMRI timeseries across voxels was extracted from these ROIs, and the same GLM described above re-fitted. Planned comparisons on the resulting parameter estimates were then tested (Table 1). OFA and FFA showed face-related activation (since they were defined this way), though EVC (defined from independent data) showed de-activation, i.e., greater activation for scrambled faces. The latter might reflect low-level differences in visual complexity, despite the phase-scrambling’s preservation of the 2D spatial power spectrum, or could reflect suppression of low-level features that are predicted by a higher-level percept (Murray & Wojciulik, 2004). All six ROIs showed significant RS for both immediate and delayed repetition, and greater RS for immediate than delayed repetition. Bilateral FFA also showed significant effects of recognition (greater activation for familiar than unfamiliar faces). Left EVC and right OFA also showed recognition effects, though would be unlikely to survive correction for the multiple comparisons performed.

**Table 1.** T-values for the effects of face perception, immediate repetition (Imm Rep), delayed repetition (Del Rep), face recognition, and their interactions on original data from 6 ROIs and on fitted data in 2-, 3- and 6-ROI networks. Note that the T-values for the “Perception” effect in OFA and FFA are biased by prior selection of the voxels in a whole-brain search; the remaining effects are unbiased since orthogonal contrasts. Positive T-values mean greater activation for faces than scrambled for the perception contrast (i.e, negative T-values mean in EVC greater activation for scrambled than intact faces), greater activation for initial than repeated presentations for the repetition contrasts (i.e, RS) and greater activation for familiar than unfamiliar faces for the recognition contrast. Second column shows mean percentage across participants of variance in fMRI timeseries explained by DCM in each ROI. *p < .05, two-tailed t-test.

| ROI | Var. Explain | Perception | Imm Rep | Del Rep | Recognition | Imm vs Delay Rep | Perception×<br>Imm Rep | Perception ×<br>Del Rep | Recognition ×<br>Imm Rep | Recognition ×<br>Del Rep |
| --- | --- | --- | --- | --- | --- | --- | --- | --- | --- | --- |
| **Original data** |  |  |  |  |  |  |  |  |  |  |
| rEVC |  | -3.08* | 3.87* | 3.75* | 0.83 | 3.40* | -1.31 | -0.27 | -1.35 | 0.44 |
| IEVC |  | -2.02 | 5.02* | 5.43* | 2.47* | 4.37* | -1.47 | -0.22 | 0.57 | 0.59 |
| rOFA |  | 13.38* | 4.06* | 3.22* | 2.60* | 3.11* | 0.70 | -0.43 | -0.84 | 0.43 |
| IOFA |  | 13.25* | 3.77* | 4.04* | 1.83 | 3.98* | 2.58* | 0.29 | 0.70 | 1.69 |
| rFFA |  | 10.94* | 6.32* | 4.54* | 5.34* | 5.04* | 1.38 | 0.05 | -0.44 | 0.46 |
| IFFA |  | 11.47* | 4.93* | 3.04* | 5.04* | 3.76* | 1.71 | -0.08 | -0.65 | 0.53 |
| **2-ROI network of right hemisphere** |  |  |  |  |  |  |  |  |  |  |
| rOFA | (29%) | 9.69* | 4.22* | 2.85* | 2.52* | 3.33* | 0.46 | 2.12*<sup>a</sup> | -0.84 | 0.60 |
| rFFA | (20%) | 9.23* | 6.51* | 5.29* | 3.68* | 6.30* | 2.94*<sup>a</sup> | 4.12*<sup>a</sup> | -0.80 | -1.16 |
| **3-ROI network of right hemisphere** |  |  |  |  |  |  |  |  |  |  |
| rEVC | (17%) | -0.75<sup>a</sup> | 2.97* | 5.86* | 0.90 | 3.71* | 1.20 | 0.65 | -0.30 | -1.13 |
| rOFA | (35%) | 8.30* | 4.90* | 3.51* | 4.22* | 3.93* | 0.50 | 1.05 | -0.28 | 0.87 |
| rFFA | (28%) | 8.46* | 5.08* | 4.31* | 3.90* | 4.43* | 0.69 | 0.85 | -1.41 | 0.03 |
| **6-ROI network of bilateral hemispheres** |  |  |  |  |  |  |  |  |  |  |
| rEVC | (16%) | 0.83<sup>a</sup> | 4.70* | 4.23* | 2.34*<sup>a</sup> | 4.19* | 0.49 | -1.51 | -0.55 | 1.01 |
| IEVC | (18%) | 0.02 | 5.62* | 4.97* | 1.42<sup>a</sup> | 5.13* | 0.77 | -1.13 | -0.42 | 1.26 |
| rOFA | (34%) | 8.03* | 5.19* | 3.41* | 4.67* | 4.05* | 0.40 | -0.15 | -0.49 | 0.83 |
| IOFA | (29%) | 8.72* | 5.37* | 3.43* | 4.87*<sup>a</sup> | 3.92* | 0.42<sup>a</sup> | -0.17 | -0.33 | 1.21 |
| rFFA | (25%) | 8.05* | 5.77* | 4.42* | 4.30* | 4.70* | 0.50 | -0.11 | -0.42 | 1.23 |
| IFFA | (22%) | 7.75* | 5.84* | 4.20* | 4.40* | 4.40* | -0.52 | -0.56 | -0.73 | 1.54 |

The only interaction reaching significance was between face perception and immediate repetition in left OFA. No other interactions between perception and repetition, or between recognition and repetition, were significant in any ROI. The lack of interactions was somewhat surprising, in that we expected RS in OFA and FFA to be greater for familiar than unfamiliar faces, and for faces than for scrambled faces (Henson & Rugg, 2003), but this may be because the repetition lags were shorter than used previously (Henson, 2016).

Given the lack of interactions, for the DCM B matrices, we only modelled the four experimental effects that significantly modulated activation in at least two ROIs. These were the main effects of “face perception” (contrast vector = [1 1 1 1 1 1 0 0 0], where order of conditions as above), “immediate repetition” (contrast vector = [0 1 0 0 1 0 0 1 0]), “delayed repetition” (contrast vector = [0 0 1 0 0 1 0 0 1]) and “face recognition” (of familiar faces; contrast vector = [1 1 1 0 0 0 0 0 0]). For the driving input in the DCM C matrix, we used the common effect of all stimuli versus inter-stimulus baseline (“all stimuli”, contrast vector = [1 1 1 1 1 1 1 1 1]).

### 2.3 Dynamic Causal Modelling (DCM)

The neural dynamics in DCM for fMRI data are represented by the first-order differential equation (Friston et al., 2003):

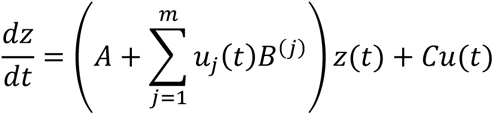

The vector ***z****(t)* represents the neural activity in each of the *n* ROIs at time *t*. The *n* × *n* matrices **A** and **B**^(j)^ are directional connectivity matrices, where the value in row *r* and column *c* represents the connection strength from ROI *c* to ROI *r* (where 0 = no connection present). The **A** matrix captures the fixed (or endogenous) connections, whereas **B**^(j)^ is a modulation on one or more of these connections due to the *j*th experimental manipulation. Each **B** matrix is multiplied by experimental input *u_j_(t)* relating to experimental effects *j=1..m* (i.e, one of the four contrasts described above). The *n* × *p* **C** matrix is the influence of one or more of those experimental inputs to one or more ROIs in the network (here *p=1*, corresponding to all stimuli versus baseline; as above). All inputs were mean-centred, so the parameters in the **A** matrix represent the average effective connectivity across conditions. All connection parameters in **A**, **B**, **C** are rate constants with units of Hz. The diagonal elements of the **A** and **B** matrices (self-connections and modulations on self-connections respectively) are always negative (inhibitory), so that the dynamics of the system settles back to baseline after stimulation. These self-connections are log scaling parameters that scale the default value of -0.5 Hz, i.e. total self-connection = -0.5 × exp(A_ii_ + B_ii_). DCM includes “shrinkage” priors on all connections, so their expected value is 0 unless the data requires otherwise.

The neural activity in each ROI is then transformed into the modelled fMRI BOLD signal using a nonlinear haemodynamic model with three main parameters (for more details, see Stephan et al., 2007). These parameters have tight empirical priors, but can differ between ROIs in order to capture different neurovascular coupling across the brain and across individuals. The combined neural and haemodynamic parameters are estimated by fitting the fMRI data from all ROIs using an iterative scheme that maximizes the free energy bound on the Bayesian model evidence, which offers a balance between explaining the data and minimizing model complexity.

In more detail, we used a recent extension of DCM to fit multi-subject data using Parametric Empirical Bayes (PEB). PEB introduced a general linear model (GLM) that encodes between-participant effects on the DCM parameters (Zeidman et al., 2019). Together, the within-participant DCMs and the group-level GLM form a Bayesian hierarchical model. This can be used in an iterative fashion to “rescue” subjects who fall into local optima, by estimating the group-average connectivity parameters, then using these parameters to form *empirical priors* for the within-participant DCMs. The DCMs are then re-estimated with these empirical priors, and the process repeats (Friston et al., 2015). We used this iterative fitting, applied to the A and B connections only, with a single covariance component to quantify between-participant variability, since preliminary analyses showed that this produced the highest free-energy approximation to the model evidence (except for the 2 ROI model, where we also applied PEB to the C connection, so as to be able to test models differing in the input ROI). We estimated the “full” PEB model (with all possible connections of interest), before applying Bayesian Model Comparison (BMC) to make inferences about sets of parameters (e.g., “self” vs “between-region”, or “forward” vs “backward” connections), by grouping PEB models into two ‘families’ (for a given hypothesis) and pooling evidence within each family.

### 2.4 DCM networks

Three sets of networks were estimated (see Figure 1), starting from a simple 2-ROI OFA + FFA network in the right hemisphere, to mimic prior analysis by Ewbank et al. (2013), and given prior evidence that the right hemisphere is particularly involved in face processing (Ishai et al., 2005; Rossion, 2018). We then added a third ROI (right EVC) to form a 3-ROI network, to allow for the important possibility that effects of repetition, face perception and/or face recognition already arise in the inputs to OFA and FFA (see Introduction). Finally, we modelled a bilateral network with 3 ROIs per hemisphere, and connections between homologous OFA and FFA, to allow for possible inter-hemisphere modulations, as proposed by Frässle et al. (2016). The 2-ROI and 3-ROI networks in the left hemisphere were also estimated and shown as supplementary results, to allow comparison with the results from the right hemisphere.

**Figure 1.**
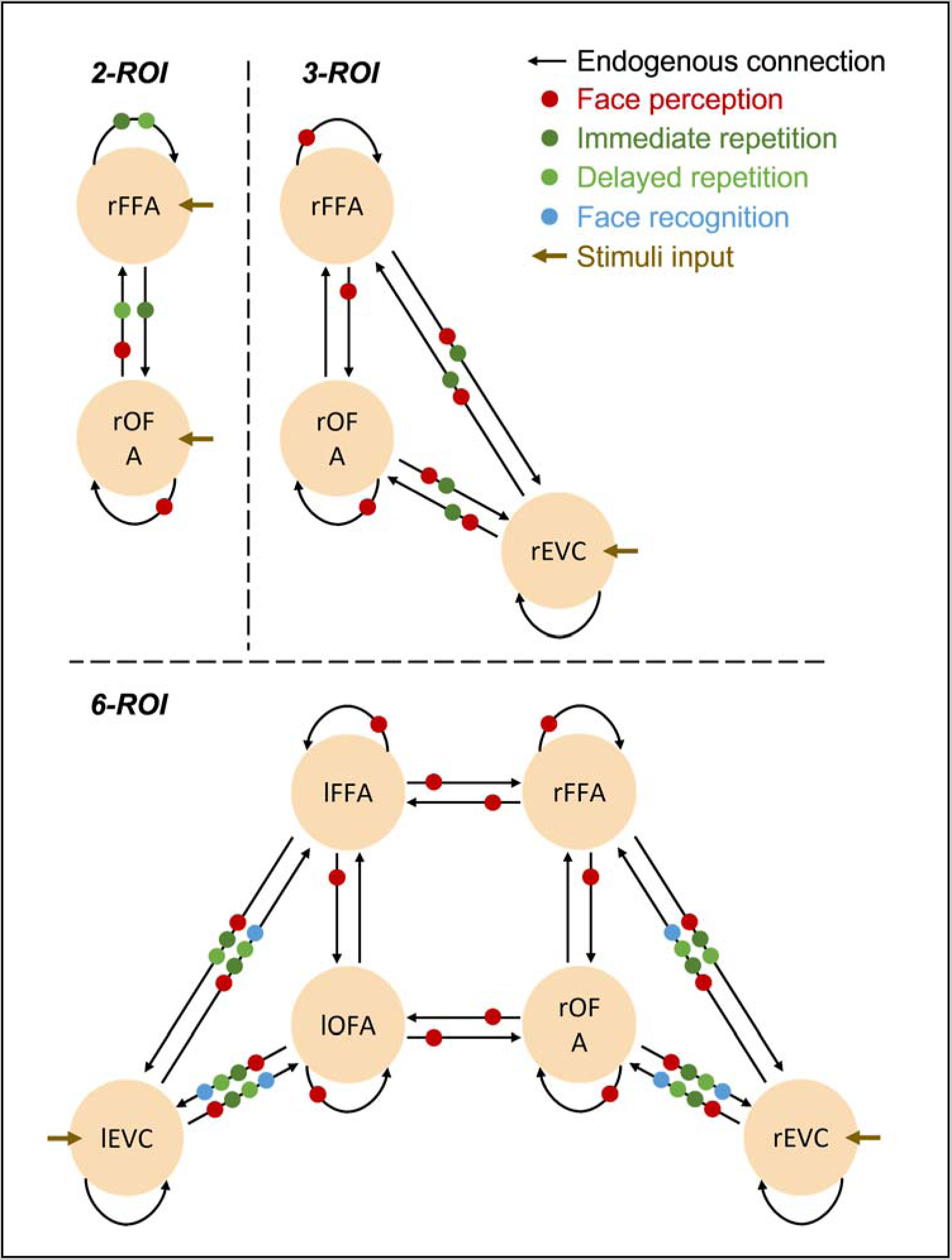
The “full” DCM structures of 2-ROI (top left), 3-ROI (top right) and 6-ROI (bottom) models. Black arrows represent endogenous (i.e., task-independent) connections (DCM “A” matrix). Colored dots represent reliable modulations (DCM “B” matrix) inferred from model comparison (see Results). Brown arrows represent driving inputs (DCM “C” matrix).

#### 2.4.1 2-ROI network

In the 2-ROI network, all possible connections were included, i.e., four endogenous connections (in a 2 × 2 **A** matrix) representing the average connectivity for OFA-self, FFA-self, OFA-to-FFA, and FFA-to-OFA. All four of these connections were allowed to be modulated by each of the four experimental effects (the four, 2 × 2 **B** matrices), i.e., face perception, immediate repetition, delayed repetition and face recognition. Importantly (compared to 3-ROI network below), the driving input (**C** matrix) for all stimuli entered into both OFA and FFA. After fitting this model to all participants using PEB, we tested a priori hypotheses using BMC.

Foremost, we tested for each of the four modulatory effects: 1) whether self-modulations are needed, by grouping all 16 models into two families based on whether a model has an OFA-self and/or an FFA-self modulation, and 2) whether any between-region modulation was needed, depending on whether the OFA-to-FFA and/or FFA-to-OFA connection was present. If evidence was found for modulation of self- or between-region modulation, further binary BMC was used to test individual self-connections and individual directions of between-region connections (e.g., OFA to FFA, or FFA to OFA).

Secondly, we tested whether the input was needed to one or both of OFA and FFA. In a strict version of the standard hierarchical model, input enters the OFA before being passed on to FFA. However, given Rossion et al’s work (Rossion, 2008; Rossion et al., 2003; see Introduction), we compared this model to models in which input was to FFA instead, or both OFA and FFA.

#### 2.4.2 3-ROI network

In the 3-ROI network, all possible nine connections were also included in the A and B matrices, but now the driving input (C matrix) was restricted to only enter through the EVC ROI, to capture the expected flow of visual information from early to later visual regions. In principle, this allowed DCM to drop (set to zero) connections from EVC to OFA or FFA, depending on whether information passes serially through OFA before reaching FFA, or whether it passes through FFA before OFA (see Introduction), or whether there are direct routes from EVC to both OFA and FFA. For the family BMC, we conducted 4 family comparisons: (1) whether self-modulation (self-OFA and/or self-FFA) was needed, (2) whether modulation between OFA and FFA (OFA-to-FFA and/or FFA-to-OFA) was needed, (3) whether “forward” modulation (EVC-to-OFA and/or EVC-to-FFA) was needed, and (4) whether “backward” modulation (OFA-to-EVC and/or FFA-to-EVC) was needed. If any modulation was found (e.g., between OFA and FFA), we further tested which direction of connectivity was modulated (e.g., OFA to FFA, or FFA to OFA).

#### 2.4.3 6-ROI network

In the 6-ROI network, the 3-ROI network for the right hemisphere was connected to the 3-ROI network for the left hemisphere, in order to account for any interhemispheric integration of face perception (Frässle et al., 2016). More specifically, homologous connections between left and right OFA and left and right FFA were modelled (no direct connections between left and right EVC were included). All connections had experimental modulations. We performed the same family-wise BMC as in the 2- and 3-ROI networks above, as well as a further family BMC to test whether interhemispheric modulation (OFA-to-OFA and FFA-to-FFA) was needed. Again, if BMC showed at least one of these connection types was needed (e.g., between left and right FFA), we went further to test which direction of connectivity was modulated (e.g., left FFA to right FFA, or vice versa).

## 3 Results

### 3.1 Univariate Results and Validation of DCM fit

Before reviewing the DCM connectivity parameters, we first validated whether the various DCMs captured significant effects in the data. First, we examined the proportion of variance in the original fMRI timeseries explained in each ROI. Averaged across participants, this ranged from 16% to 35% across ROIs (Table 1), which is generally good for DCM, given the typical amount of noise in fMRI data (though tended to be lower in EVC, most likely because this ROI was defined independent of the current data).

However, DCM could fit a reasonable percentage of the variance in the fMRI timeseries by assuming that every stimulus produced an evoked response (versus interstimulus baseline) of equal amplitude, i.e., without necessarily reproducing the significant differences between conditions (stimulus-types) that was found in the data. To check the latter, we extracted the timeseries predicted by DCM, fit the same GLM that was applied to the data timeseries (i.e., with a separate regressor for each of the nine conditions) and performed the same T-contrasts on the resulting parameter estimates that were performed in Table 1. Note that these parameter estimates are not a perfect reflection of DCM’s predictions, because they assume a fixed HRF (the canonical HRF used in the GLM), whereas DCM allows the HRF to differ across participants and ROIs, but the results should be similar nonetheless.

Figure 2 presents the parameter estimates from the GLM fit to the original data, and from the same GLM fit to the timeseries generated by DCM for each of the 2-, 3- and 6-ROI networks; Table 1 lists the T-values of planned comparisons on these parameters from the DCM fit (cf. the T-values of original data, see the top part of Table 1). In general, the relative pattern of significant differences across conditions was reproduced in left and right OFA and FFA by all models, i.e., effects of perception, recognition, immediate repetition, delayed repetition, and difference between immediate and delayed repetition. However, the 3-ROI and 6-ROI networks did not reproduce the greater activation for scrambled than intact faces in right EVC (despite the presence of backward connections from OFA/FFA to EVC). Also, the effects of perception, immediate repetition, delayed repetition, and difference between immediate and delayed repetition, were reproduced in all ROIs in left-hemisphere networks, but a significant effect of recognition was also produced in lOFA (Supplementary Table 1).

**Figure 2.**
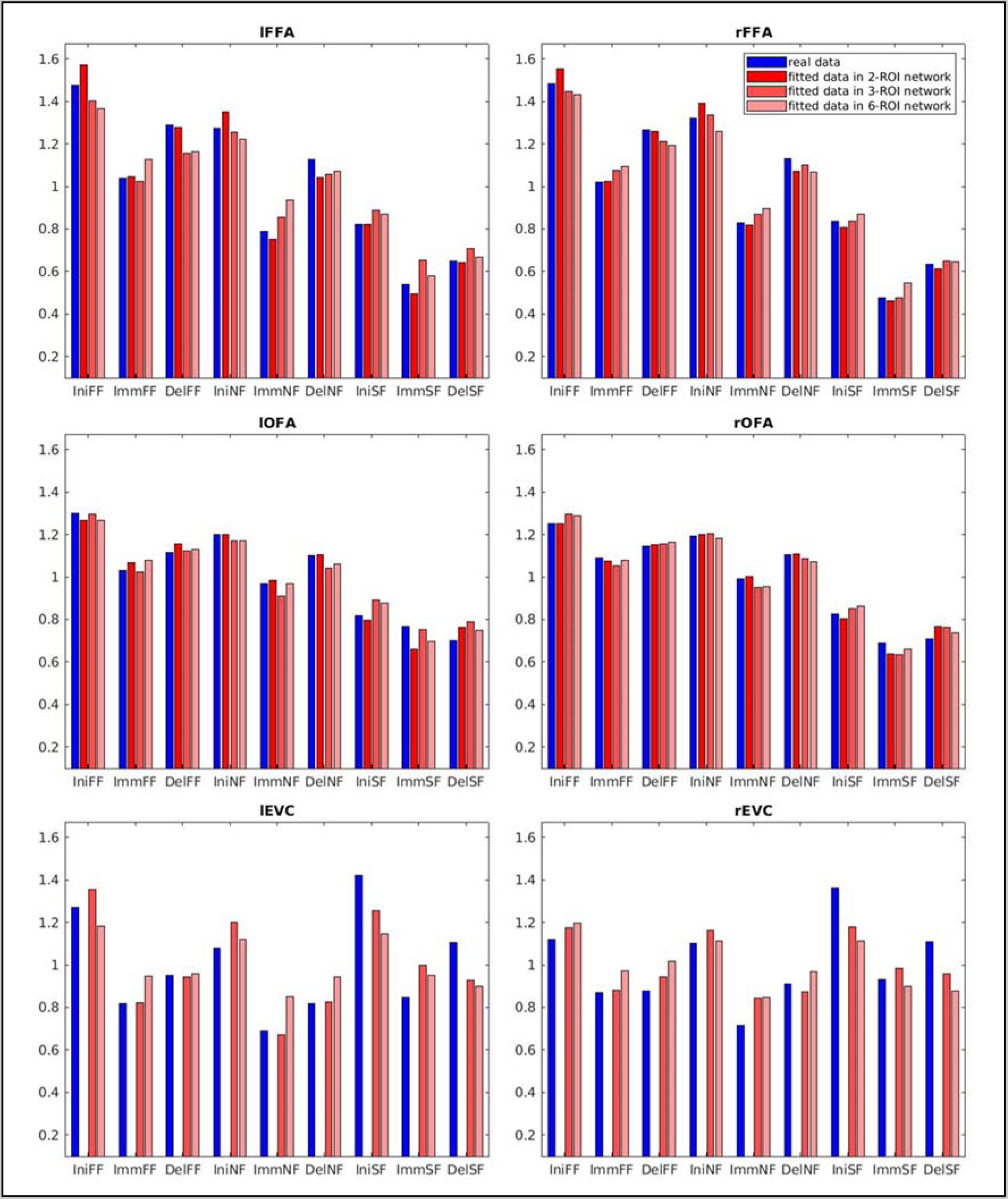
Mean GLM parameter estimates from original timeseries (blue) and from timeseries reconstructed by DCM for 2ROI, 3ROI and 6ROI networks (red shades), for each ROI (panel) and condition (groups on x-axis). To allow for different scaling factors, the parameter estimates were re-scaled to have same mean over all conditions. The nine conditions are initial familiar face (IniFF), immediately repeated familiar face (ImmFF), delayed repeated familiar face (DelFF), initial unfamiliar face (IniNF), immediately repeated unfamiliar face (ImmNF), delayed repeated unfamiliar face (DelNF), initial scrambled face (IniSF), immediately repeated scrambled face (ImmSF), and delayed repeated scrambled face (DelSF).

### 3.2 Connectivity Results

#### 3.2.1 2-ROI network

To address the theories in the Introduction about whether specific types of modulatory connection were affected by face perception, repetition and recognition, BMC was performed on pairs of model families with versus without certain connection-types.

There was strong evidence (> 95% probability) that face perception modulated both self-connections of rOFA and/or rFFA, and between-region connections between rOFA and rFFA (Table 2). Indeed, follow-up tests showed that modulations by faces was needed for the rOFA self-connection and the connection from rOFA to rFFA.

**Table 2.**
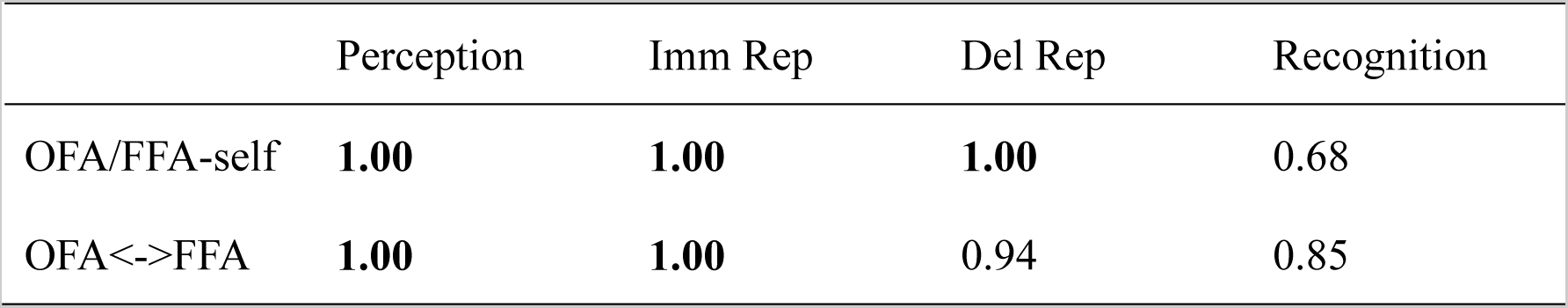
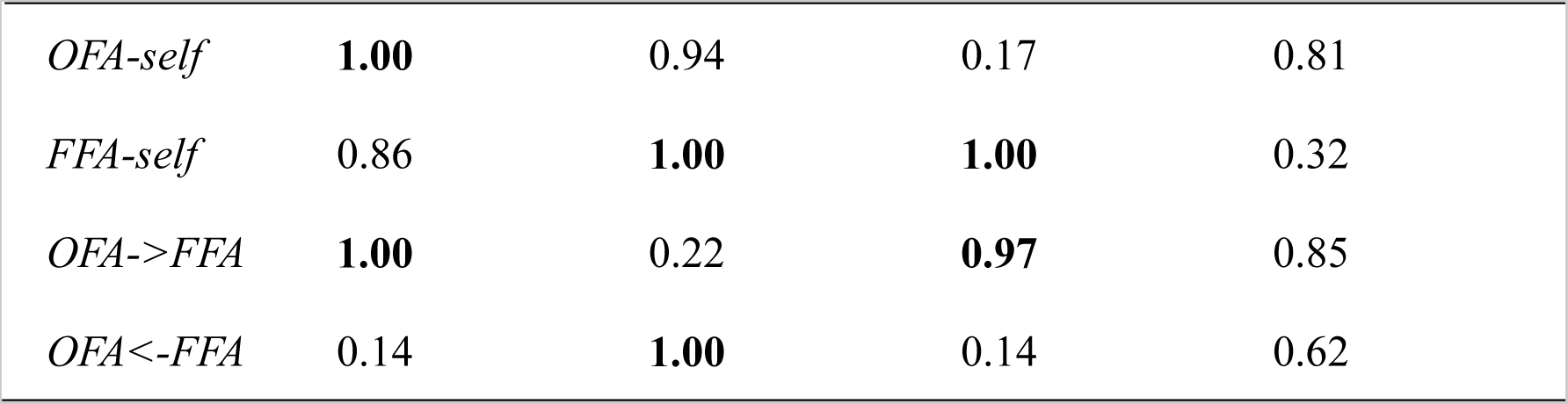
Posterior probabilities for BMC of two families with versus without various types of connection (rows) for each experimental effect (columns) for the 2-ROI, right hemisphere network. Values greater than 0.95 are taken as strong evidence (shown in bold emphasis).

There was strong evidence that immediate and delayed repetition modulated self-connections of rOFA and/or rFFA, with follow-up tests showing that modulation of rFFA was critical. There was also evidence that the two types of repetition modulated between-region connections differently, with immediate repetition modulating the connection from rFFA to rOFA, and delayed repetition modulating the connection from rOFA to rFFA.

There was insufficient evidence to identify which specific connections were modulated by face recognition in this right hemisphere network.

The results for the left hemisphere DCM were similar (Supplementary Table 2), in that face perception affected the lOFA self-connection and the lOFA->lFFA connection, that repetition affected the lFFA self-connection, and that delayed repetition modulated the lOFA- >lFFA connection (though modulation by immediate repetition could no longer be attributed to the lFFA->lOFA connection). Unlike the right hemisphere DCM, face recognition now modulated the lOFA->lFFA connection (suggesting that recognition effects were stronger in left hemisphere – see later).

We also tested whether inputs were needed to just one or both ROIs (C parameters). There was compelling evidence (posterior probability close to 1.00) that stimulus-dependent input (regardless of stimulus type) was needed to both ROIs, consistent with Rossion et al. (2008), and further justifying consideration of the 3-ROI network below.

The above results suggest that repetition primarily affects both self-connections and between-region connections. However, this assumes that the input to both ROIs not already modulated by repetition. Thus in the next model, we added a third ROI, EVC, connected to both OFA and FFA, which allowed us to ask whether repetition (and perception/recognition) modulated input from EVC to OFA and/or FFA.

#### 3.2.2 3-ROI network

There was evidence that all the connections were modulated by face perception, except from rOFA to rFFA (Table 3).^1^ This included a direct connection from rEVC to rFFA (as well as from rEVC to rOFA), consistent with Rossion’s non-hierarchical view. The modulations of connections from rOFA and rFFA back to rEVC were probably needed to explain the greater activation for scrambled vs intact faces in rEVC (see Univariate Results).

**Table 3.** Posterior probabilities for BMC of two families with versus without various types of connection (rows) for each experimental effect (columns) for the 3-ROI, right hemisphere network. Values greater than 0.95 are taken as strong evidence (shown in bold emphasis).

|  | Perception | Imm Rep | Del Rep | Recognition |
| --- | --- | --- | --- | --- |
| OFA/FFA-self | **1.00** | 0.39 | 0.49 | 0.59 |
| OFA<->FFA | **1.00** | 0.51 | 0.30 | 0.44 |
| EVC->OFA/FFA | **1.00** | **1.00** | 0.19 | 0.32 |
| EVC<-OFA/FFA | **1.00** | **1.00** | 0.18 | 0.20 |
| *OFA-self* | **0.99** | 0.52 | 0.55 | 0.35 |
| *FFA-self* | **1.00** | 0.35 | 0.36 | 0.69 |
| *OFA->FFA* | 0.70 | 0.25 | 0.25 | 0.26 |
| *OFA<-FFA* | **1.00** | 0.68 | 0.41 | 0.59 |
| *EVC->OFA* | **1.00** | **0.99** | 0.23 | 0.41 |
| *EVC->FFA* | **1.00** | **1.00** | 0.23 | 0.31 |
| *OFA->EVC* | **1.00** | **1.00** | 0.22 | 0.26 |
| *FFA->EVC* | **1.00** | **1.00** | 0.22 | 0.23 |

For immediate repetition however, modulations could no longer be attributed to self-connections of rOFA and rFFA, nor the direct connection between them. Rather, it was the connections from, and to, EVC that showed evidence of modulation. For delayed repetition, on the other hand, there were no longer evidence that could uniquely identify the source of modulation. Like for the 2-ROI right hemisphere network, there was not sufficient evidence to localise modulations by face recognition either.

The results for the left hemisphere DCM (Supplementary Table 3) showed similar modulations by face perception, except that the lOFA self-connection and lEVC->lFFA connection no longer showed sufficient evidence. However, family comparison could not uniquely localise modulation by immediate or delayed repetition. There was however evidence that face recognition modulated the lOFA self-connection and lFFA->lEVC connection, again suggesting left lateralisation of this connectivity changes associated with face recognition.

In summary, the effects of face perception on self-connections and connections between rOFA and rFFA in the 3-ROI network are consistent with the 2-ROI network, but additionally suggest that faces already start to differ from scrambled faces in the input to OFA and FFA (a scenario that was not possible to test in the 2-ROI network). This pattern is more consistent with a non-hierarchical view, where face information is present in a direct input to FFA (at least in the right hemisphere), rather than conventional hierarchical view in which face information in FFA only comes via the OFA.

On the other hand, the effects of repetition, at least immediate repetition, are different from the 2-ROI network, in that whereas immediate repetition modulated both self-connections and between-region connections in the 2-ROI network, it now modulated only the connections between EVC-OFA and EVC-FFA in the 3-ROI network (at least in the right hemisphere). This is again because the 2-ROI network does not have the capability to explain repetition effects as arising earlier in the visual pathway. This result from the 3-ROI network instead favours theories that entail changes in between-region connectivity (e.g., synchronization or predictive coding) rather than local changes within a region RS (e.g., fatigue or sharpening). Further implications of these results are considered in the Discussion, after comparing with results from the final 6-ROI network.

#### 3.2.3 6-ROI network

The 6-ROI network combines the right and left 3-ROI networks, with additional “inter-hemispheric” connections between rOFA and lOFA, and rFFA and lFFA (but not EVC).

For face perception, there was evidence that all connections were modulated, except for the OFA->FFA connections, in both directions (Table 5), like for the 3-ROI networks above. There was also evidence that faces modulated inter-hemispheric connections for OFA and FFA in both directions.

**Table 5.** Posterior probabilities for BMC of two families with versus without various types of connection (rows) for each experimental effect (columns) for the 6-ROI, bilateral network. Values greater than 0.95 are taken as strong evidence (shown in bold emphasis).

|  | Perception | Imm Rep | Del Rep | Recognition |
| --- | --- | --- | --- | --- |
| *OFA/FFA-self, l/r* | **1.00** | 0.26 | 0.54 | 0.09 |
| *OFA<->FFA, l/r* | **0.98** | 0.13 | 0.59 | 0.04 |
| *EVC->OFA/FFA, l/r* | **1.00** | **1.00** | **1.00** | **1.00** |
| *EVC<-OFA/FFA, l/r* | **1.00** | **1.00** | **1.00** | 0.90 |
| *l<->r, OFA/FFA* | **1.00** | 0.03 | 0.04 | 0.06 |
| *OFA-self, l/r* | **1.00** | 0.44 | 0.75 | 0.15 |
| *FFA-self, l/r* | **1.00** | 0.09 | 0.11 | 0.11 |
| *EVC->OFA, l/r* | **1.00** | **1.00** | **1.00** | **1.00** |
| *EVC->FFA, l/r* | **1.00** | **1.00** | **1.00** | **0.98** |
| *OFA->EVC, l/r* | **1.00** | **1.00** | **1.00** | **0.96** |
| *FFA->EVC, l/r* | **1.00** | **1.00** | **1.00** | 0.04 |
| *OFA->FFA, l/r* | 0.19 | 0.02 | 0.03 | 0.06 |
| *FFA->OFA, l/r* | **0.99** | 0.29 | 0.81 | 0.06 |
| *r->l, OFA/FFA* | **1.00** | 0.07 | 0.02 | 0.10 |
| *l->r, OFA/FFA* | **1.00** | 0.20 | 0.10 | 0.05 |

For immediate repetition, there was again compelling evidence for modulation of connections from, and to, EVC, but no evidence of modulation of self-connections and direct connections between rOFA and rFFA, as in the 3-ROI, right hemisphere network. There was no evidence of modulation of inter-hemispheric connections. Unlike the 3-ROI, right hemisphere network, there was also evidence that delayed repetition modulated the same connections as immediate repetition, i.e., from EVC to OFA/FFA and vice versa. This might reflect the doubling in the amount of data being fit.

Finally, for face recognition, there was evidence for modulation of EVC-to-OFA and EVC-to-FFA modulations, as well as OFA-to-EVC. This pattern is unlike the 3-ROI networks, but may reflect the pooling across both hemispheres (see supplementary Table 3).

## 4 Discussion

In this study, we applied Dynamical Causal Modelling (DCM) with Parametric Empirical Bayes (PEB) on a publically available fMRI dataset in order to estimate the effective connectivity within networks including the left and/or right face-sensitive regions of occipital face area (OFA) and fusiform face area (FFA), plus input from early visual cortex (EVC), in response to initial and repeated presentations of familiar faces, unfamiliar faces and scrambled faces. We applied DCM to unilateral 2-ROI and 3-ROI networks, as well as a bilateral 6-ROI network, but focus on those effects that were consistent across these networks.

### 4.1 Face repetition effects

Our main interest concerned the effects of immediate and delayed repetition of stimuli, specifically whether the well-documented repetition suppression (RS) in OFA and FFA is best explained by local changes (self-connections in DCM), as predicted by fatigue and sharpening theories, or by between-ROI connections, as predicted by synchronization and predictive coding theories (see Introduction). When using a simple 2-ROI network of right OFA and right FFA, as in Ewbank et al. (2013), we found evidence that both immediate and delayed repetition modulated the FFA self-connection and connections between OFA and FFA (and similarly for the two homologous regions in the left hemisphere). This is not consistent with the findings of Ewbank et al., who found that repetition affected only the connection from OFA to FFA (when the face image was the same size, as here). This discrepancy could reflect several factors, including the present use of a randomized rather than blocked design, where a randomized design reduces the influence of expectation of repetition (Henson, 2016).

More importantly however, the 2-ROI network does not allow repetition to modulate the input to OFA and/or FFA. In other words, the 2-ROI model considered by Ewbank et al (2013) does not allow RS to arise earlier in the visual processing pathway, i.e., in the inputs to OFA and/or FFA. To accommodate this, we also fit a 3-ROI network in which a third ROI, early visual cortex (EVC), was connected to both OFA and FFA. For the right hemisphere, we now found that immediate repetition modulated both “forward” and “backward” connections between rEVC and rOFA/FFA, but there was no longer evidence that it modulated the direct connections between rOFA and rFFA, or the self-connections of rFFA (or rOFA). Thus contrary to the 2-ROI architecture, the 3-ROI architecture, by allowing repetition to modulate the input to OFA and FFA, instead favoured synchronization or predictive coding accounts of RS, at least for immediate repetition.

Our last step was to test the modulation of repetition in the 6-ROI network, which allowed additional interhemispheric modulation. Like for the 3-ROI, right hemisphere network, we found that immediate repetition modulated the connections between EVC and OFA/FFA, but not direct connections between OFA and FFA, nor self-connections of OFA/FFA. The same pattern was now also found for delayed repetition. Thus, the consistent findings of between-region modulations across 2-, 3- and 6-ROI networks suggests that RS is caused by interactions between regions, as predicted by synchronization and predictive coding models. It should be noted however that synaptic depression (one possible neural mechanism of fatigue model) could explain reduced effective connectivity between regions if it is the synapses from EVC to OFA/FFA that are depressed by prior processing – i.e., mapping of Fatigue to local effects not simple.

Finally, note that these repetition effects were averaged across familiar, unfamiliar and scrambled faces, because we did not find significant interactions between repetition and stimulus-type in the univariate activation of the six ROIs (except in lOFA, but this would not survive correction for multiple comparisons across ROIs). For example, one might have expected greater RS in OFA and FFA for faces than scrambled faces. This lack of interactions was true for both immediate and delayed repetition, despite the greater RS overall for immediate repetition (Table 1). The lack of interactions was surprising, because we have found such interactions in previous work (e.g., Henson et al., 2000), though these tended to use larger lags between initial and repeated presentations, and lag may modulate repetition effects (Henson, 2016).

### 4.2 Face perception effects

When we compared whether the input was to one or both ROIs in the 2-ROI network, we found strong evidence that input to both ROIs was needed. This is consistent with imaging evidence from patients with OFA lesions reported by Rossion et al. (2008), who still showed FFA activation. The 3-ROI network provided further support for this, with evidence that connections from EVC to both OFA and FFA were modulated by faces. This suggests that face information is already extracted in the transformations from EVC to OFA and FFA, contrary to the “standard” hierarchical model that assumes that input to FFA only arises from OFA.

Several previous DCM studies have examined the face modulations among EVC, OFA and FFA. Consistent with our results, Lohse et al. (2016) found face modulation on “forward” connections from rEVC to rOFA and to rFFA, while Furl et al. (2015) found face modulation on the connection from rEVC to rOFA only. Frässle et al.’s (2016) study was the only one to include bilateral EVC, OFA and FFA. Their results revealed face modulation on connections from EVC to OFA, and between right and left OFA, like in our 6-ROI network, but also from OFA to FFA, unlike in our 3- and 6-ROI networks. However, the latter is most likely because they did not allow any direct connections (and hence modulations) between EVC and FFA. In addition, none of these studies allowed modulations on self-connections. Our study estimated all possible DCM connections and modulations among EVC, OFA and FFA, and showed that “forward” (EVC to OFA), “backward” (OFA to EVC) and inter-hemispheric modulations were needed for face perception in our data, but not direct connections between OFA and FFA.

While our 6-ROI results were generally comparable with our 3-ROI results, the striking differences between our 2-ROI and 3-ROI results highlight the importance of the model architecture when testing hypotheses with DCM. While our results show that the addition of an EVC region has important effects on the conclusions one draws, it is possible that the results would change again if other regions were added, like anterior temporal lobes, amygdala (Xiu et al., 2015), or in particular superior temporal sulcus (STS), which is well-known to have face-responsive neurons (Furl et al., 2015; Kessler et al., 2021; Lohse et al., 2016). Indeed, using an exhaustive data-driven approach called “Group Iterative Multiple Model Estimation” (Gates & Molenaar, 2012), Elbich et al. (2019) found that connections from STS to OFA/FFA were also modulated by faces. Though STS did not show significant face-related activation that surpassed our corrected threshold, which is why we did not include it here, future work could add STS to DCM networks like the ones here.

There are other differences between prior studies that may also affect which connections are modulated by face processing, such as the specific stimuli contrasted with faces (e.g., phase-scrambled faces, as here, versus non-face stimuli like objects or cars, Furl et al., 2015; Lohse et al., 2016) or the type of design (e.g., randomized versus blocked, Frässle et al., 2016; Furl et al., 2015; Lohse et al., 2016), which can affect top-down expectancies for a certain type of stimulus. The role of these factors could be explicitly tested in future empirical studies.

### 4.3 Face recognition effects

When analysing the right hemisphere networks, we were not able to uniquely attribute face recognition effects to specific connection-types. Note that this does not mean that DCM could not reproduce the greater activations to familiar faces that was found in many of the ROIs (including right hemisphere; Figure 2); it just means that the timeseries related to familiar faces did not allow inference about which type of connection would uniquely reproduce the distinct parts of that timeseries (compared to other stimulus types). In other words, it is possible that changes in self, forward or backward connections could equally well explain the distinct part of the ROI timeseries related to face recognition. When analysing the left hemisphere networks, on the other hand, it appeared that the lOFA self-connection was modulated by face recognition.

However, when analysing the bilateral 6-ROI network, a different result emerged, with face recognition modulating forward connections from EVC to OFA and FFA (averaged across hemispheres), as well as from OFA to EVC. Given these quite different results across the various networks, we remain cautious about interpreting connectivity changes associated with recognition of familiar faces, particularly since such recognition may also involve interactions with more anterior regions like anterior temporal lobes (ATL) and orbitofrontal cortex (OFC) (Fairhall & Ishai, 2007), and possibly even left lateral prefrontal regions associated with covert naming of known faces. Future DCM models could investigate whether the greater activation to familiar faces in the present ROIs reflect top-down feedback from regions “higher” on the visual processing pathway (e.g., using better matched stimuli, cognitive tasks that explicitly require face identification, and fMRI sequences optimised to handle signal drop-out in OFC and ATL).

### 4.4 Hemispheric Differences

Most studies examined connectivity among regions in the right hemisphere because of the hypothesized specialization of right hemisphere for face processing (Kanwisher et al., 1997). However, Frässle et al. (2016) included OFA and FFA in both hemispheres, and claimed that the right lateralization of the activation pattern was due to an interhemispheric modulation from left to right OFA. Furthermore, because their face images were presented in either the right or left visual field, they found evidence of modulation of connections from EVC to OFA in both hemispheres by both the visual field and the presence of faces. Though faces were presented centrally in our data, our 6-ROI network also found modulation by faces on both interhemispheric connections and EVC-to-OFA connections in both hemispheres, supporting the claim that face processing is not specific to the right hemisphere.

### 4.5 Limitations and future directions

There are several methodological limitations of this study. First, any DCM analysis evaluates models defined within a certain architecture (determined by the number of ROIs and connections allowed between them). This means there may be other more probable models comprising different regions and connections. In this study, we focused on two regions showing significant experimental effects of face and repetition, OFA and FFA, and their likely input region, EVC, and allowed for some variations in architecture by focusing on results that were consistent across 3 different networks (2-ROI, 3-ROI and 6-ROI networks). More generally, one could use lots of ROIs and evaluate whether a method like Bayesian model reduction (BMR) or Group Iterative Multiple Model Estimation will reveal the most parsimonious set of connections between them. However, when we tried BMR here, the results were difficult to interpret, at least when more than two ROIs, most likely because of the high co-dependency between connection parameters in such fully-connected and recurrent networks, and so we resorted to more hypothesis-driven BMC to focus on specific connection types.

The second limitation is that, while both synchronization and predictive coding models are consistent with the general concept of repetition affecting connectivity between regions, the direction of this effective connectivity remains unclear. In predictive coding for example, repetition both improves top-down predictions and reduces bottom-up prediction errors, and it is unclear how either of these relate precisely to forward and backward fMRI connectivity. One way to address this issue is to apply DCM to EEG and/or MEG data. The much richer dynamics in EEG/MEG evoked responses allows fitting of more complex neurophysiological models, e.g., “canonical microcircuit” model (Bastos et al., 2012), in which forward and backward connections operate with different timescales. Furthermore, predictive coding does not rule out within-region (self-connection) modulation too, particularly in more complex implementations that allow for interactions between cells within different layers of cortex within the same ROI. Future high-resolution fMRI, e.g., at 7T, might allow separate modelling of cortical layers. Predictive coding theory would predict that repetition causes reduced activity in superficial pyramidal cells in the supragranular layer, and changes in its connectivity with inhibitory interneurons in other layers.

### 4.6 Conclusion

To conclude, we used DCM to examine the effective connectivity among face-selective regions during face repetition, face perception and face recognition. The simplest conclusion about repetition suppression is that it reflects more than local changes within a region, so fatigue or sharpening models are not sufficient; rather, the repetition-related modulation of between-region connections is consistent with synchronization and/or predictive coding models of repetition suppression (Ewbank & Henson, 2012). While the effective connectivity associated with recognition remains unclear, a consistent finding regarding face perception is that it includes modulation of connections direct from EVC to FFA, without needing modulation from OFA to FFA, which supports recent suggestions for a non-hierarchical view of the “core” face network in the posterior ventral stream.

## Acknowledgements

S-M.L. was supported by the Study Abroad Program of Ministry of Science and Technology, Taiwan (108-2917-I-006-009); R.T. was supported by a British Academy Postdoctoral Fellowship (SUAI/028 RG94188). P.Z. was supported by core funding from Wellcome awarded to the Wellcome Centre for Human Neuroimaging (203147/Z/16/Z). P.S.Y. was supported by core funding from SRK Medicare Pvt. Ltd., India. R.H. was supported by UK Medical Research Council programme grant (SUAG/086 G116768). For the purpose of open access, the authors have applied a Creative Commons Attribution (CC BY) licence to any Author Accepted Manuscript version arising from this submission.

## Footnotes

1 Because we were only interested in the univariate results for OFA and FFA, we did not distinguish families by self-connections for EVC, even though the DCM model allowed modulation of EVC self-connections.

## References

Avidan, G., Hasson, U., Hendler, T., Zohary, E., & Malach, R. (2002). Analysis of the Neuronal Selectivity Underlying Low fMRI Signals. Current Biology, 12(12), 964–972. https://doi.org/10.1016/S0960-9822(02)00872-2

Babo-Rebelo, M., Puce, A., Bullock, D., Hugueville, L., Pestilli, F., Adam, C., Lehongre, K., Lambrecq, V., Dinkelacker, V., & George, N. (2022). Visual Information Routes in the Posterior Dorsal and Ventral Face Network Studied with Intracranial Neurophysiology and White Matter Tract Endpoints. Cerebral Cortex, 32(2), 342–366. https://doi.org/10.1093/cercor/bhab212

Bastos, A. M., Usrey, W. M., Adams, R. A., Mangun, G. R., Fries, P., & Friston, K. J. (2012). Canonical Microcircuits for Predictive Coding. Neuron, 76(4), 695–711. https://doi.org/10.1016/j.neuron.2012.10.038

Damasio, A. R., Tranel, D., & Damasio, H. (1990). Face Agnosia and the Neural Substrates of Memory. Annual Review of Neuroscience, 13(1), 89–109. https://doi.org/10.1146/annurev.ne.13.030190.000513

Desimone, R. (1996). Neural mechanisms for visual memory and their role in attention. Proceedings of the National Academy of Sciences, 93(24), 13494–13499. https://doi.org/10.1073/pnas.93.24.13494

Egner, T., Monti, J. M., & Summerfield, C. (2010). Expectation and Surprise Determine Neural Population Responses in the Ventral Visual Stream. Journal of Neuroscience, 30(49), 16601–16608. https://doi.org/10.1523/JNEUROSCI.2770-10.2010

Elbich, D. B., Molenaar, P. C. M., & Scherf, K. S. (2019). Evaluating the organizational structure and specificity of network topology within the face processing system. Human Brain Mapping, 40(9), 2581–2595. https://doi.org/10.1002/hbm.24546

Epstein, R. A., Parker, W. E., & Feiler, A. M. (2008). Two Kinds of fMRI Repetition Suppression? Evidence for Dissociable Neural Mechanisms. Journal of Neurophysiology, 99(6), 2877–2886. https://doi.org/10.1152/jn.90376.2008

Ewbank, M. P., & Henson, R. N. (2012). Explaining away repetition effects via predictive coding. Cognitive Neuroscience, 3(3–4), 239–240. https://doi.org/10.1080/17588928.2012.689960

Ewbank, M. P., Henson, R. N., Rowe, J. B., Stoyanova, R. S., & Calder, A. J. (2013). Different Neural Mechanisms within Occipitotemporal Cortex Underlie Repetition Suppression across Same and Different-Size Faces. Cerebral Cortex, 23(5), 1073–1084. https://doi.org/10.1093/cercor/bhs070

Ewbank, M. P., Lawson, R. P., Henson, R. N., Rowe, J. B., Passamonti, L., & Calder, A. J. (2011). Changes in “Top-Down” Connectivity Underlie Repetition Suppression in the Ventral Visual Pathway. Journal of Neuroscience, 31(15), 5635–5642. https://doi.org/10.1523/JNEUROSCI.5013-10.2011

Fairhall, S. L., & Ishai, A. (2007). Effective Connectivity within the Distributed Cortical Network for Face Perception. Cerebral Cortex, 17(10), 2400–2406. https://doi.org/10.1093/cercor/bhl148

Frässle, S., Paulus, F. M., Krach, S., Schweinberger, S. R., Stephan, K. E., & Jansen, A. (2016). Mechanisms of hemispheric lateralization: Asymmetric interhemispheric recruitment in the face perception network. NeuroImage, 124, 977–988. https://doi.org/10.1016/j.neuroimage.2015.09.055

Friston, K. J. (2005). A theory of cortical responses. Philosophical Transactions of the Royal Society B: Biological Sciences, 360(1456), 815–836. https://doi.org/10.1098/rstb.2005.1622

Friston, K. J., Harrison, L., & Penny, W. (2003). Dynamic causal modelling. NeuroImage, 19(4), 1273–1302. https://doi.org/10.1016/S1053-8119(03)00202-7

Friston, K. J., Zeidman, P., & Litvak, V. (2015). Empirical Bayes for DCM: A Group Inversion Scheme. Frontiers in Systems Neuroscience, 9. https://www.frontiersin.org/article/10.3389/fnsys.2015.00164

Furl, N., Henson, R. N., Friston, K. J., & Calder, A. J. (2015). Network Interactions Explain Sensitivity to Dynamic Faces in the Superior Temporal Sulcus. Cerebral Cortex, 25(9), 2876–2882. https://doi.org/10.1093/cercor/bhu083

Gainotti, G., & Marra, C. (2011). Differential Contribution of Right and Left Temporo-Occipital and Anterior Temporal Lesions to Face Recognition Disorders. Frontiers in Human Neuroscience, 5. https://doi.org/10.3389/fnhum.2011.00055

Gates, K. M., & Molenaar, P. C. M. (2012). Group search algorithm recovers effective connectivity maps for individuals in homogeneous and heterogeneous samples. NeuroImage, 63(1), 310–319. https://doi.org/10.1016/j.neuroimage.2012.06.026

Gentile, F., Ales, J., & Rossion, B. (2017). Being BOLD: The neural dynamics of face perception. Human Brain Mapping, 38(1), 120–139. https://doi.org/10.1002/hbm.23348

Ghuman, A. S., Bar, M., Dobbins, I. G., & Schnyer, D. M. (2008). The effects of priming on frontal-temporal communication. Proceedings of the National Academy of Sciences, 105(24), 8405–8409. https://doi.org/10.1073/pnas.0710674105

Gotts, S. J. (2016). Incremental learning of perceptual and conceptual representations and the puzzle of neural repetition suppression. Psychonomic Bulletin & Review, 23(4), 1055–1071. https://doi.org/10.3758/s13423-015-0855-y

Gotts, S. J., Chow, C. C., & Martin, A. (2012). Repetition priming and repetition suppression: Multiple mechanisms in need of testing. Cognitive Neuroscience, 3(3–4), 250–259. https://doi.org/10.1080/17588928.2012.697054

Grill-Spector, K., Henson, R. N., & Martin, A. (2006). Repetition and the brain: Neural models of stimulus-specific effects. Trends in Cognitive Sciences, 10(1), 14–23. https://doi.org/10.1016/j.tics.2005.11.006

Grill-Spector, K., Kushnir, T., Edelman, S., Avidan, G., Itzchak, Y., & Malach, R. (1999). Differential Processing of Objects under Various Viewing Conditions in the Human Lateral Occipital Complex. Neuron, 24(1), 187–203. https://doi.org/10.1016/S0896-6273(00)80832-6

Grill-Spector, K., & Malach, R. (2001). fMR-adaptation: A tool for studying the functional properties of human cortical neurons. Acta Psychologica, 107(1), 293–321. https://doi.org/10.1016/S0001-6918(01)00019-1

Grotheer, M., & Kovács, G. (2015). The relationship between stimulus repetitions and fulfilled expectations. Neuropsychologia, 67, 175–182. https://doi.org/10.1016/j.neuropsychologia.2014.12.017

Haxby, J. V., Hoffman, E. A., & Gobbini, M. I. (2000). The distributed human neural system for face perception. Trends in Cognitive Sciences, 4(6), 223–233. https://doi.org/10.1016/S1364-6613(00)01482-0

Henson, R. N. (2016). Repetition suppression to faces in the fusiform face area: A personal and dynamic journey. Cortex, 80, 174–184. https://doi.org/10.1016/j.cortex.2015.09.012

Henson, R. N., Goshen-Gottstein, Y., Ganel, T., Otten, L. J., Quayle, A., & Rugg, M. D. (2003). Electrophysiological and Haemodynamic Correlates of Face Perception, Recognition and Priming. Cerebral Cortex, 13(7), 793–805. https://doi.org/10.1093/cercor/13.7.793

Henson, R. N., & Mouchlianitis, E. (2007). Effect of spatial attention on stimulus-specific haemodynamic repetition effects. NeuroImage, 35(3), 1317–1329. https://doi.org/10.1016/j.neuroimage.2007.01.019

Henson, R. N., & Rugg, M. D. (2003). Neural response suppression, haemodynamic repetition effects, and behavioural priming. Neuropsychologia, 41(3), 263–270. https://doi.org/10.1016/S0028-3932(02)00159-8

Henson, R. N., Shallice, T., & Dolan, R. (2000). Neuroimaging Evidence for Dissociable Forms of Repetition Priming. Science, 287(5456), 1269–1272. https://doi.org/10.1126/science.287.5456.1269

Ishai, A., Schmidt, C. F., & Boesiger, P. (2005). Face perception is mediated by a distributed cortical network. Brain Research Bulletin, 67(1), 87–93. https://doi.org/10.1016/j.brainresbull.2005.05.027

Jiang, X., Bradley, E., Rini, R. A., Zeffiro, T., VanMeter, J., & Riesenhuber, M. (2007). Categorization Training Results in Shape- and Category-Selective Human Neural Plasticity. Neuron, 53(6), 891–903. https://doi.org/10.1016/j.neuron.2007.02.015

Jiang, X., Rosen, E., Zeffiro, T., VanMeter, J., Blanz, V., & Riesenhuber, M. (2006). Evaluation of a Shape-Based Model of Human Face Discrimination Using fMRI and Behavioral Techniques. Neuron, 50(1), 159–172. https://doi.org/10.1016/j.neuron.2006.03.012

Kanwisher, N., McDermott, J., & Chun, M. M. (1997). The Fusiform Face Area: A Module in Human Extrastriate Cortex Specialized for Face Perception. Journal of Neuroscience, 17(11), 4302–4311. https://doi.org/10.1523/JNEUROSCI.17-11-04302.1997

Kessler, R., Rusch, K. M., Wende, K. C., Schuster, V., & Jansen, A. (2021). Revisiting the effective connectivity within the distributed cortical network for face perception. Neuroimage: Reports, 1(4), 100045. https://doi.org/10.1016/j.ynirp.2021.100045

Larsson, J., Solomon, S. G., & Kohn, A. (2016). FMRI adaptation revisited. Cortex, 80, 154–160. https://doi.org/10.1016/j.cortex.2015.10.026

Li, L., Miller, E. K., & Desimone, R. (1993). The representation of stimulus familiarity in anterior inferior temporal cortex. Journal of Neurophysiology, 69(6), 1918–1929. https://doi.org/10.1152/jn.1993.69.6.1918

Lohse, M., Garrido, L., Driver, J., Dolan, R. J., Duchaine, B. C., & Furl, N. (2016). Effective Connectivity from Early Visual Cortex to Posterior Occipitotemporal Face Areas Supports Face Selectivity and Predicts Developmental Prosopagnosia. Journal of Neuroscience, 36(13), 3821–3828. https://doi.org/10.1523/JNEUROSCI.3621-15.2016

McMahon, D. B. T., & Olson, C. R. (2007). Repetition Suppression in Monkey Inferotemporal Cortex: Relation to Behavioral Priming. Journal of Neurophysiology, 97(5), 3532–3543. https://doi.org/10.1152/jn.01042.2006

Murray, S. O., & Wojciulik, E. (2004). Attention increases neural selectivity in the human lateral occipital complex. Nature Neuroscience, 7(1), 70–74. https://doi.org/10.1038/nn1161

Rossion, B. (2008). Constraining the cortical face network by neuroimaging studies of acquired prosopagnosia. NeuroImage, 40(2), 423–426. https://doi.org/10.1016/j.neuroimage.2007.10.047

Rossion, B. (2018). Damasio’s error – Prosopagnosia with intact within-category object recognition. Journal of Neuropsychology, 12(3), 357–388. https://doi.org/10.1111/jnp.12162

Rossion, B., Caldara, R., Seghier, M., Schuller, A., Lazeyras, F., & Mayer, E. (2003). A network of occipito-temporal face-sensitive areas besides the right middle fusiform gyrus is necessary for normal face processing. Brain, 126(11), 2381–2395. https://doi.org/10.1093/brain/awg241

Rowe, J. B., Hughes, L. E., Barker, R. A., & Owen, A. M. (2010). Dynamic causal modelling of effective connectivity from fMRI: Are results reproducible and sensitive to Parkinson’s disease and its treatment? NeuroImage, 52(3), 1015–1026. https://doi.org/10.1016/j.neuroimage.2009.12.080

Steeves, J., Dricot, L., Goltz, H. C., Sorger, B., Peters, J., Milner, A. D., Goodale, M. A., Goebel, R., & Rossion, B. (2009). Abnormal face identity coding in the middle fusiform gyrus of two brain-damaged prosopagnosic patients. Neuropsychologia, 47(12), 2584–2592. https://doi.org/10.1016/j.neuropsychologia.2009.05.005

Stephan, K. E., Weiskopf, N., Drysdale, P. M., Robinson, P. A., & Friston, K. J. (2007). Comparing hemodynamic models with DCM. NeuroImage, 38(3), 387–401. https://doi.org/10.1016/j.neuroimage.2007.07.040

Summerfield, C., Trittschuh, E. H., Monti, J. M., Mesulam, M.-M., & Egner, T. (2008). Neural repetition suppression reflects fulfilled perceptual expectations. Nature Neuroscience, 11(9), 1004–1006. https://doi.org/10.1038/nn.2163

Wakeman, D. G., & Henson, R. N. (2015). A multi-subject, multi-modal human neuroimaging dataset. Scientific Data, 2, 150001. https://doi.org/10.1038/sdata.2015.1

Wiggs, C. L., & Martin, A. (1998). Properties and mechanisms of perceptual priming. Current Opinion in Neurobiology, 8(2), 227–233. https://doi.org/10.1016/S0959-4388(98)80144-X

Xiu, D., Geiger, M. J., & Klaver, P. (2015). Emotional face expression modulates occipital-frontal effective connectivity during memory formation in a bottom-up fashion. Frontiers in Behavioral Neuroscience, 9. https://doi.org/10.3389/fnbeh.2015.00090

Zeidman, P., Jafarian, A., Seghier, M. L., Litvak, V., Cagnan, H., Price, C. J., & Friston, K. J. (2019). A guide to group effective connectivity analysis, part 2: Second level analysis with PEB. NeuroImage, 200, 12–25. https://doi.org/10.1016/j.neuroimage.2019.06.032

