## Supplemental Figures and Tables for "Effects of Face Repetition on Ventral Visual Stream Connectivity using Dynamic Causal Modelling of fMRI data"

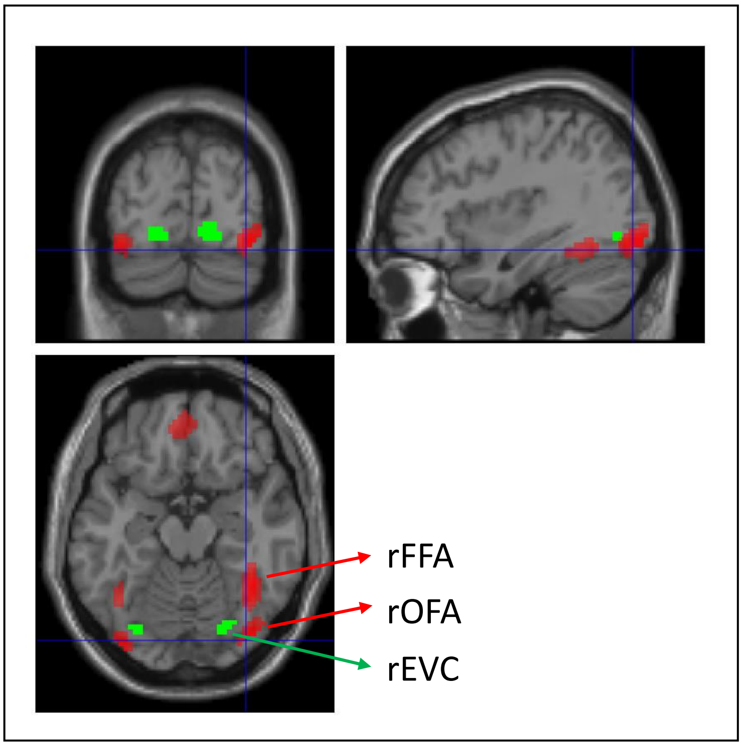


*Supplementary Figure 1. The locations of ROIs. Red clusters, namely bilateral occipital face area (OFA) and fusiform face area (FFA), were defined by the contrast of faces > scrambled, uncorrected p < .05, in a group-level analysis. Green clusters, namely bilateral early visual cortex (EVC), were defined by the contrast of left > right and right > left hemifield input from Henson & Mouchlianitis (2007). Note that the location and size of ROIs was variable across participants (see Methods).*

*Supplementary Table 1. T-values of main effects of face perception, immediate repetition (Imm Rep), delayed repetition (Del Rep), face recognition, and interactions of Imm Rep × Del Rep, Perception × Imm Rep, Perception × Del Rep, Recognition × Imm Rep, Recognition × Del Rep on fitted data in 2- and 3-ROI networks of left hemisphere. The mean across participants of the proportion of original fMRI timeseries variance explained by the model in each ROI is shown in brackets. *p < .05, two-tailed t-test. [a]The effects are inconsistent (one is significant but another one is insignificant) in real (Table 1) and fitted data.*

| | **Perception** | **Imm Rep** | **Del Rep** | **Recognition** | **Imm vs Del Rep** | **Perception × Imm Rep** | **Perception × Del Rep** | **Recognition × Imm Rep** | **Recognition × Del Rep** |
|---|---|---|---|---|---|---|---|---|---|
| | **2-ROI network of left hemisphere** | | | | | | | | |
| lOFA (.24) | 9.55* | 5.06* | 3.62* | 2.95*[a] | 4.56* | 2.05[a] | 2.84*[a] | -0.78 | 0.47 |
| lFFA (.14) | 6.85* | 6.38* | 4.95* | 3.09* | 6.53* | 3.60*[a] | 3.91*[a] | -1.29 | -0.72 |
| | **3-ROI network of left hemisphere** | | | | | | | | |
| lEVC (.17) | -0.98 | 5.21* | 5.95* | 2.49* | 6.05* | 2.68*[a] | 0.95 | 0.03 | 0.49 |
| lOFA (.29) | 6.24* | 4.32* | 3.57* | 3.29*[a] | 4.17* | 1.86[a] | 1.86 | 0.15 | 2.05 |
| lFFA (.23) | 5.82* | 4.42* | 3.86* | 2.87* | 4.16* | 1.79 | 0.84 | -0.23 | 0.89 |

*Supplementary Table 2. Posterior probabilities for family-wise BMC across several types of connection (rows) for each experimental effect (columns) for the 2-ROI model of left hemisphere. Values greater than 0.95 are taken as strong evidence (shown in bold emphasis). Rounded to 2 decimal places.*

| | Perception | Imm Rep | Del Rep | Recognition |
|---|---|---|---|---|
| OFA/FFA-self | **1.00** | **1.00** | **1.00** | 0.35 |
| OFA<->FFA | **1.00** | 0.48 | **0.96** | **0.99** |
| *OFA-self* | **1.00** | 0.38 | 0.16 | 0.48 |
| *FFA-self* | 0.81 | **1.00** | **1.00** | 0.27 |
| *OFA->FFA* | **1.00** | 0.14 | **0.99** | **0.99** |
| *OFA<-FFA* | 0.21 | 0.69 | 0.13 | 0.74 |

Supplementary Table 3. Posterior probabilities for family-wise BMC across several types of connection (rows) for each experimental effect (columns) for the 3-ROI model of left hemisphere. Values greater than 0.95 are taken as strong evidence (shown in bold emphasis). Rounded to 2 decimal places.

| | **Perception** | **Imm Rep** | **Del Rep** | **Recognition** |
|---|---|---|---|---|
| OFA/FFA-self | **1.00** | 0.71 | 0.45 | **1.00** |
| OFA<->FFA | **1.00** | 0.36 | 0.19 | 0.78 |
| EVC->OFA/FFA | **1.00** | 0.92 | 0.21 | 0.28 |
| EVC<-OFA/FFA | **1.00** | 0.38 | 0.77 | **0.96** |
| *OFA-self* | 0.27 | 0.29 | 0.58 | **0.97** |
| *FFA-self* | **1.00** | 0.84 | 0.29 | 0.88 |
| *OFA->FFA* | 0.36 | 0.42 | 0.22 | 0.24 |
| *OFA<-FFA* | **1.00** | 0.35 | 0.25 | 0.88 |
| *EVC->OFA* | **1.00** | 0.92 | 0.29 | 0.37 |
| *EVC->FFA* | 0.88 | 0.79 | 0.22 | 0.27 |
| *OFA->EVC* | **1.00** | 0.53 | 0.63 | 0.39 |
| *FFA->EVC* | **1.00** | 0.25 | 0.84 | **0.98** |
